# Discovery of a AhR flavonoid agonist that counter-regulates ACE2 expression in rodent models of inflammation and attenuates ACE2-SARS-CoV2 interaction *in vitro*

**DOI:** 10.1101/2021.02.24.432203

**Authors:** Michele Biagioli, Silvia Marchianò, Rosalinda Roselli, Cristina Di Giorgio, Rachele Bellini, Martina Bordoni, Anna Gidari, Samuele Sabbatini, Daniela Francisci, Bianca Fiorillo, Bruno Catalanotti, Eleonora Distrutti, Adriana Carino, Angela Zampella, Gabriele Costantino, Stefano Fiorucci

**Affiliations:** Dipartimento di Medicina e Chirurgia, Università di Perugia; Perugia, Italy; University of Naples Federico II, Department of Pharmacy, Naples, Italy; SC di Gastroenterologia ed Epatologia, Azienda Ospedaliera di Perugia; Perugia, Italy; Department of Food and Drugs, University of Parma, Parma. Italy

**Keywords:** Pelargonidin, TNF-α, Ahr, ACE2, SARS-CoV-2

## Abstract

The severe acute respiratory syndrome (SARS)-CoV-2, a newly emerged coronavirus first identified in 2019, is the pathogenetic agent od Corona Virus Induced Disease (COVID)19. The virus enters the human cells after binding to the angiotensin converting enzyme (ACE) 2 receptor in target tissues. ACE2 expression is induced in response to inflammation. The colon expression of ACE2 is upregulated in patients with inflammatory bowel disease (IBD), highlighting a potential risk of intestinal inflammation in promoting viral entry in the human body. Because mechanisms that regulate ACE2 expression in the intestine are poorly understood and there is a need of anti-SARS-CoV2 therapies, we have settled to investigate whether natural flavonoids might regulate the expression of ACE2 in intestinal models of inflammation. The results of these studies demonstrated that pelargonidin, a natural flavonoid bind and activates the Aryl hydrocarbon Receptor (AhR) in vitro and reverses intestinal inflammation caused by chronic exposure to high fat diet or to the intestinal braking-barrier agent DSS in a AhR-dependent manner. In these two models, development of colon inflammation associated with upregulation of ACE2 mRNA expression. Colon levels of ACE2 mRNA were directly correlated with TNFα mRNA levels. In contrast to ACE2 the angiotensin 1-7 receptor MAS was downregulated in the inflamed tissues. Molecular docking studies suggested that pelargonidin binds a fatty acid binding pocket on the receptor binding domain of SARS-CoV2 Spike protein. *In vitro* studies demonstrated that pelargonidin significantly reduces the binding of SARS-CoV2 Spike protein to ACE2 and reduces the SARS-CoV2 replication in a concentration-dependent manner. In summary, we have provided evidence that a natural flavonoid might hold potential in reducing intestinal inflammation and ACE2 induction in the inflamed colon in a AhR-dependent manner.

## Introduction

The coronavirus disease 2019 (COVID-19) is a respiratory tract infection caused severe acute respiratory syndrome (SARS)-CoV-2, a newly emerged coronavirus first identified in the city of Wuhan in China in December 2019 [1]. The SARS-CoV2 virus entry into host cells is mediated by interaction between the spike (S) glycoprotein, that assembly into an homotrimeric complex protruding from the viral surface, with the angiotensin-converting enzyme 2 (ACE2) [2]. The spike glycoprotein comprises two functional subunits: the S1 subunit contains a specific motif, the receptor-binding domain (RBD), that is responsible for binding to ACE2, while the S2 subunit is essential for fusion of the viral and cellular membranes [2] Thus, targeting the interaction of spike’s RBD with ACE2 might hold potential for the treatment of COVID 19. Accordingly, several agents that target the SARS-CoV2/ACE2 interaction, including several monoclonal antibodies, have been identified and tested clinically. Following a similar approach, it has been suggested that modulation of ACE2 expression could also be of interest in reducing SARS-CoV2 entry in target cells [2–7].

The expression of ACE2 vary greatly among human tissues. In the normal lung, ACE2 mRNA is mainly expressed by type I and II alveolar (AT1 and AT2) epithelial cells and endothelial cells [8, 9]. However, it has been shown that the expression of ACE2 in the lung increases in response to inflammation or exposure to interferon IFN-γ [10]. In normal individual, ACE2 mRNA and protein are highly expressed in the gastrointestinal system, in particular in the duodenum, jejunum, ileum, and colon, although the functional role of this peptidase in intestinal epithelial cells remains portly elucidated [11]. Furthermore, similarly to the lung, the intestinal expression of ACE2 is modulated in response to intestinal inflammation in inflammatory bowel disease (IBD) in a phenotype dependent manner[12]. More specifically, while expression of Ace2 is reduced in the ileum of patients with active Crohn disease, an elevated colonic expression of ACE2 mRNA has been reported in ulcerative colitis patients with active inflammation and severe disease [13–15]. Since the colonic ACE2 levels normalize after anti-cytokine therapy and the anti-cytokine therapy associates with reduced morbidity of IBD patients from COVID19, these data establish a relation between the levels of colonic ACE2 expression and beneficial effects exerts by anti-cytokine therapies. Moreover, the evaluation of SECURE-IBD registry (available http://www.covidibd.org.) suggest that patients with IBD are under-represented in those diagnosed with COVID-19 compared to the general populations [16]. Despite these data raise interest on ACE2 as potential target in COVID19 and IBD, the mechanisms involved in ACE2 regulation in the intestine are only partially understood. Further on, it is unclear whether ACE2 expression could be modulated with dietary factors that might have also utility in reducing inflammation.

The aryl hydrocarbon receptor (AhR) is a ligand-activated nuclear receptor that belongs to the basic helix-loop-helix (bHLH)/PerARNT-Sim (PAS) superfamily. Although AhR was originally described as a xenobiotics sensor, it is now well established that dietary/intestinal microbiota metabolites represent the physiological ligands [17]. Similarly, to other nuclear receptors, in the absence of a ligand, AhR resides in cytoplasm as a component of a chaperone complex [18]. The AhR is widely expressed by cells of innate/adaptive immunity [19] but it is also expressed at high levels by intestinal epithelial cells where it is essential for maintaining the integrity of the intestinal barrier and therefore for regulating the inflammatory state of the gastrointestinal tract. [20, 21].

Pelargonidin is a water soluble anthocyanidin that is widely diffuse in nature as glycosylated derivatives and that is responsible for the red/orange colours of berries such as raspberries and strawberries, as well as blueberries and blackberries [22]. While, similarly to other anthocyanins, pelargonidin is thought to be beneficial for human health [23–25], controversies exist over the doses needed to reach these effects, along with poor systemic bioavailability and unclear mechanism(s) of action. In a previous study we have shown that the natural pelargonidin function as AhR ligand in vitro and attenuates intestinal inflammation in AhR-dependent manner [26]. In the present study we have investigated the potential of the natural pelargonidin in regulating ACE2 expression in colon in models of intestinal inflammation caused by exposure of wild type and AhR^−/−^ mice to a high caloric intake and intestinal irritants. The results of these studies show that not only a dietary agent might regulate the expression of the ACE2 in the colon but also inhibit the interaction of SARS-CoV2’s spike protein with ACE2, providing a potentially useful clues in the prevention and treatment of COVID19.

## Material and Methods

### Transactivation Assay

For AhR-mediated transactivation, HepG2 cells were plated at 7.5 × 10^4^ cells/well in a 24-well plate. Cells were transiently transfected with 200 ng of the reporter vector pLightAhRE, 100 ng of pCMVXL4AhR, and 100 ng of pGL4.70 (Promega, Madison, WI, USA), a vector encoding the human Renilla luciferase gene. At 24 h post-transfection, cells were stimulated 18 h with pelargonidin 10 and 50 μM. Dose-response curves were performed in HepG2 cells transfected with AhR, as described above, and then treated with increasing concentrations of pelargonidin (1, 2.5 5, 10, 12.5, 25, 50, 75, 100 μM).

At 18 h post-stimulation, cellular lysates were assayed for luciferase and Renilla activities using the Dual-Luciferase Reporter assay system (E1980, Promega, Madison, WI, USA). Luminescence was measured using Glomax 20/20 luminometer (Promega, Madison, WI, USA). Luciferase activities (RLU) were normalized with Renilla activities (RRU).

### Mouse spleen macrophages purification

Spleens were collected from AhR^+/+^ and AhR^−/−^ mice. Spleens were homogenized and then red blood cell lysis was performed using a hypotonic solution. After cell count, the cell pellet was resuspended at a concentration of 10^8^/ mL, to proceed to the positive selection of Cd11b^+^ cells. In particularly, macrophages were purified using Cd11b microbeads (130-049-601) by Miltenyi Biotec, according to the manufacturer’s instructions. Subsequently the selected macrophages have been plated at a concentration of 500.000/mL and were activated with LPS (5 ng / mL, Sigma-Aldrich L2880) in combination with IFN-γ (20ng / mL, eBioscience, San Diego, CA, USA) for 16 hours alone or in addition to Pelargonidin (20 μM) for gene expression profile analysis.

### Animals

All animal care and experimental procedures complied with the guidelines of Animal Care and were approved by Use Committee of the University of Perugia and by the Italian Minister of Health and Istituto Superiore di Sanità (Italy) and was in agreement with the European guidelines for use of experimental animals (permissions n. 583/2017-PR and 1126/2016-PR). The general health conditions of the animals were monitored daily by the veterinarian in the animal facility. The study protocol caused minor suffering. However, mice with an hight grade of pain were eutanized. AhR knock out mice (Ahr^−/−^) on C57BL/6 background and their C57BL/6 congenic littermates wild type (Ahr^+/+^) were originally supplied by Charles River (Wilmington, MA, USA). Colonies were housed under controlled temperature (22 °C) and photoperiods of 12:12-h light/dark cycle in restricted access area and abled to acclimate to these conditions for at least 7 days before inclusion in an experiment.

To reproduce a mouse model of Acute Colitis a 2,4,6-trinitrobenzenesulfonic acid (Sigma Chemical Co, St Louis, MO) (TNBS)-colitis model is used as previously described [27]. Briefly, according to this 8–10 weeks old male mice C57BL/6J wild-type and Ahr^+/+^ and Ahr^−/−^ on C57BL/6J genetic background were administrated. Briefly, mice were fasted for 12 h overnight (Day -1). The next day (Day 0), mice were sedated by administration of Zoletil at a dose of 50 mg/Kg of body weight. Then a 3.5 F = 11.55 mm catheter was inserted into the colon such that it was up to 4 cm from the anus and 1 mg of TNBS in 50% ethanol administered via the catheter into the colon lumen using a 1 mL syringe (injection volume of 100 μL). Control mice received 50% ethanol alone. Pelargonidins were administered by oral gavage (1, 5 or 10 mg/kg) daily from day 0 to the 4 day, that is the day of sacrifice. The severity of colitis was scored daily for each mouse by assessing body weight, the fecal occult blood, and stool consistency. Each parameter was scored from 0 to 4 and the sum represents the Colitis Disease Activity Index (CDAI). The scoring system was as follows: Percent of body weight loss: None = 0; 1–5% = 1; 5–10% = 2; 10–20% = 3; >20% = 4. Stool consistency: Normal = 0; soft but still formed = 1; very soft = 2; diarrhea = 3; liquid stools that stick to the anus or anal occlusion = 4. Fecal blood: None = 0; visible in the stool = 2; severe bleeding with fresh blood around the anus and very present in the stool = 4. On day 4, surviving mice were sacrificed, blood samples collected by facial vein, and the colon excised, measured length and weight, and evaluated for macroscopic damage.

To investigate the effects of Pelargonidins in a mouse model of NASH, 10-12 weeks old C57BL/6 male mice, Ahr wild type (Ahr^+/+^) and their congenic littermates AhR null mice (Ahr^−/−^) were fed a high fat diet containing 59 KJ% fat plus 1% cholesterol, w/o sugar (ssniff® EF R/M acc. D12330 mod. 22.7 ME/Kg) and fructose in drinking water (42 g/L) for 8 weeks as previously described [28]. Food intake was valued as the difference of weight between the provided and the remnant amount of food at 2-day intervals. The food was provided as pressed pellets and the residual spillage was not considered. After 8 days, Ahr^+/+^ and Ahr^−/−^ HFD-F mice were randomized to receive HFD-F alone or plus Pelargonidin (5 mg/Kg/die) by oral gavage for remaining 7 weeks. On the sacrifice day, fed mice were deeply anesthetized with Zoletil at a dose of 50 mg/Kg and sacrificed before 12 AM. Thus, blood, liver, small intestine, colon, spleen, epididymal adipose tissue (eWAT), brown adipose tissue (BAT) and gastrocnemius were collected. The abdominal circumference (AC) (immediately anterior to the forefoot), body weight, and body length (nose-to-anus or nose–anus length), were measured in anaesthetized mice at the day of sacrifice. The body weight and body length were used to calculate the Body mass index (BMI) (=body weight (g)/length^2^ (cm^2^)).

### Intestinal Permeability

Intestinal permeability test was performed using fluorescein isothiocyanate conjugated dextran (FITC-dextran) (Sigma-Aldrich, St. Louis, MO, USA, catalog number: FD4). FITC-dextran was dissolved in PBS at a concentration of 100 mg/mL. The evening before the test day mice were fasted overnight. In the next morning, after weighing each mouse, FITC was administered to each one at a dose of 44 mg/100 g of body weight by oral gavage. After 4 hours, the mice were anesthetized by inhalation of isoflurane and 300-400 μl of blood collected from the facial vein in microtubes containing EDTA. Blood was stored in the dark. Once blood has been collected from all the mice, microtubes were processed to separate the serum. For analysis, serum was diluted with an equal volume of PBS. The concentration of FITC in serum was determined by spectrophoto fluorometry with an excitation of 485 nm and an emission wavelength of 528 nm using as standard serially diluted FITC-dextran (0, 125, 250, 500, 1,000, 2,000, 4,000, 6,000, 8,000 ng/ml). Serum from mice not administered with FITC-dextran was used to determine the background.

### Histology

Colon sample (2-3 cm up anus) were first fixed in 10% formalin, embedded in paraffin, cut into 5-μm-thick sections and then stained with Hematoxylin/Eosin (H&E) for histopathological analysis. The histological score of the colon was assessed as previously described by U. Erben et al [29]. This score evaluated the level of tissue inflammation in relation to the extension of the inflammatory cell infiltrate (mild severity if the cellular infiltrate is only present in the mucosa, moderate if they are involved mucosa and submucosa, marked if the infiltration is transmural), and epithelial changes of the intestinal mucosal architecture like erosions and ulcerations.

### Quantitative Real-Time PCR analysis

RNA was extracted from mouse colon and spleen macrophages using respectively TRIzol reagent (Invitrogen) and Direct-zol™ RNA MiniPrep w/Zymo-Spin™ IIC Columns (Zymo Research, Irvine, CA, USA). 1 μg of RNA from each sample was reverse transcribed using FastGene Scriptase Basic Kit (Nippon Genetics Europe) in a 20-μL reaction volume; 50 ng of cDNA was amplified in a 20-μL solution containing 200 nM of each primer and 10 μL of SYBR Select Master Mix (Thermo Fisher Scientific, Waltham, MA, USA). All reactions were performed in triplicate using the following thermal cycling conditions: 3 min at 95 °C, followed by 40 cycles of 95 °C for 15 s, 56 °C for 20 s, and 72 °C for 30 s, using a StepOnePlus system (Applied Biosystems, Foster City, CA, USA). The relative mRNA expression was calculated according to the ΔCt method. Primers were designed using the software PRIMER3 (http://frodo.wi.mit.edu/primer3/), using data published in the NCBI database. The primer used were as following (forward and reverse): *mGapdh* (for CTGAGTATGTCGTGGAGTCTAC; rev GTTGGTGGTGCAGGATGCATTG), *mTnfα* (for CCACCACGCTCTTCTGTCTA; rev AGGGTCTGGGCCATAGAACT), *mIl-6* (for CTTCACAAGTCGGAGGCTTA; rev TTCTGCAAGTGCATCATCGT), mTgfβ (for TTGCTTCAGCTCCACAGAGA; rev TGGTTGTAGAGGGCAAGGAC), *mAce2* (for AGATGGCCGGAAAGTTGTCT; rev GGGCTGTCAAGAAGTTGTCC), *mMas* (for CTGCTGACAGCCATCAGTGT; rev ACAGAAGGGCACAGACGAAT), *mIl1β* (for GCTGAAAGCTCTCCACCTCA; rev AGGCCACAGGTATTTTGTCG).

### Cell culture

Caco-2 cells, a human intestinal epithelial cell line (Sigma-Aldrich) was grown at 37 °C in D-MEM containing 10% FBS, 1% l-glutamine, and 1% penicillin/streptomycin. Cells were regularly passaged to maintain exponential growth. Caco-2 cells were classically activated with TNF-α 100 ng/ml for 24 h alone or in combination with pelargonidin (5, 10, and 20 μM) for the analysis of expression of several genes.

### ACE2/SARS-CoV-2 Spike Inhibitor Screening Assay Kit

Pelargonidin at different concentrations (1-10-20-50 μM) was tested using the ACE2: SARS-CoV-2 Spike Inhibitor Screening Assay Kit (BPS Bioscience Cat. number #79936) according to the manufacturer’s instructions. Briefly, we have thawed ACE2 protein on ice and diluted to 1 μg/ml in PBS. We have used 50 μL of ACE solution to coat a 96-well nickel-coated plate (1 hour). The plate was washed 3 times and incubated for 10 min with a Blocking Buffer. Next, 10 μL of inhibitor solution containing the pelargoidin was added to wells and incubated for 1 h at room temperature with slow shaking. For the “Positive Control” and “Blank,” 10 μL of inhibitor buffer (5% DMSO solution) were used. After the incubation (1 hour), SARS-CoV-2 Spike (RBD)-Fc was thawed on ice and diluted to 0.25 ng/μL (~5 nM) in Assay Buffer 1; 5 nM Spike protein was added to each well, except to the blank. The reaction was incubated for 1 h at room temperature, with slow shaking. After 3 washes and incubation with a Blocking Buffer (10 min), we have treated the plate with an Anti-mouse-Fc-HRP and the plate was incubated for 1 h at room temperature with slow shaking. Finally, HRP substrate was added to the plate to produce chemiluminescence, which was measured using FluoStar Omega microplate reader. To confirm the validity of the assay used in this study, remnants of plasma samples used to test levels of anti-SARS CoV2 IgG in post COVID-19 patients were used. The original samples were collected at the blood bank of Azienda Ospedaliera of Perugia from post COVID-19 donors who participate to a program of plasma biobanking. The program’s protocol included the quantitative analysis of the anti-SARS-CoV-2 IgG antibodies directed against the subunits (S1) and (S2) of the virus spike protein. An informed consent was obtained by each donor for the use of the plasma sample remnants and the protocol was approved by the Ethical Committee of the University of Perugia: authorization n. 61843 (July 13,2020).

### Virus Isolation and cell cultures

The experiments were performed in Biological Safety Level 3 (BSL-3) virology laboratory of “Santa Maria della Misericordia” Hospital, Perugia, Italy. The SARS-CoV-2 strain was isolated as previously described [30]. For viral propagation, Vero E6 cell line was maintained in Eagle minimal essential medium (MEM) supplemented with 10% foetal bovine serum (FBS) and 100 U/mL penicillin-streptomycin solution. After isolation, SARS-CoV-2 titer was determined by Median Tissue Culture Infectious Dose (TCID50) endpoint dilution and stock aliquots were stored at −80°C. The stock virus titer was 4.22 ×107 TCID50/mL.

### SARS-CoV-2 yield reduction assay

Vero E6 cells (20,000 cells/well) were seeded in 96-well clear flat-bottom plates and incubated at 37°C with 5% CO2 for 24 h. After incubation, pelargonidin at concentration of 20, 50 and 100 μM, and remdesivir 10μM (Veklury®, Gilead, US) all diluted in complete medium were added to each well and then, cells were infected with SARS-CoV-2 (50 or 100 TCID50/well). Negative controls (pelargonidin or remdesivir alone), infected positive controls (SARS-CoV-2 alone) and mock-infected cells were included in each plate. Plates were incubated at 37°C with 5% CO2 for 48 h. After the incubation, supernatants of 5 technical replicates were pooled and stored at −80°C for further analysis.

### Plaque-reduction assays

Supernatants viral titer was determined by plaque assay as previously described [31], with some modifications. Vero E6 cells (600,000 cells/well) were seeded in a 6-well plate and incubated at 37°C with 5% CO2 for 24 h. After incubation, the medium was removed and cells were infected with 500 μl of ten-fold serial dilution of supernatant previously obtained, rocking the plates every fifteen minutes. In the meanwhile, the overlay medium (complete medium with agar 0.1%) was prepared and maintained in a 50°C water bath. After 1 h of infection, the overlay medium (2 mL) was poured into each well and the plates incubated for 3 days. Finally, the overlay was discarded, cells were fixed for 30 min with 4% formalin and stained with 0.5% crystal violet. Viral titer was determined as plaque-forming units per mL, considering wells with plaques ranging from 2 to 100. For each pool of supernatants, plaque-reduction assay was performed in duplicate.

### Computational studies

Two cryo-electron microscopy structures of SARS-CoV2 were employed for docking calculations, in the central β-sheet core and in the flavonoids binding pocket (PDB ID: 6VSB) [32], while for dockings in the fatty acids (FA) pocket we used the (PDB ID 6ZB5) [33]. The receptor was treated with the Protein Preparation [34] tool implemented in Maestro ver. 11.8. (Schrödinger Release 2019-1: Maestro, New York). The 3D structure of pelargonidin was built using the Graphical User Graphical User Interface (GUI) of Maestro ver. 11.8. (Schrödinger Release 2019-1: Maestro, New York). The protonation state of pelargonidin at pH 7.4 in water has been calculated using the Epik module [35]. Finally, the compound was then minimized using the OPLS 2005 force field through 2500 iteration steps of the Polak-Ribiere Conjugate Gradient (PRCG) [36] algorithm.

The docking procedure was realized with the Glide software package (Glide, version 7.1. New York, NY: Schrödinger, LLC, 2019) using the Standard Precision (SP) algorithm of the GlideScore function [37] and the OPLS 2005 force field [38]. A grid box of 20 × 20 × 20 Å for SARS-CoV2 receptor centered on the putative binding pocket was created. A total amount of 100 poses was generated. Docking conformations of pelargonidin were then clustered based on their atomic RMSD and five clusters were obtained. Among them, only the conformation included in the most populated cluster with both the Glide Emodel and GlideScoredocking lowest-energy value was considered.

## Results

### Activity of the Natural Compounds toward AhR

We have first investigated whether the natural pelargonidin exerted agonistic effects on AhR. The activity of pelargonidin was evaluated in a luciferase reporter assay using HepG2 cells, transiently transfected with a AhR reporter gene cloned upstream to the luciferase. HepG2 were incubated with 5 nM TCDD, an AhR agonist as a control, or vehicle (0.1% v/v DMSO) in the presence of the pelargonidin (10-50 μM) for 18 h. As shown in Figure 1A, pelargonidin was found effective in transactivating the AhR; furthermore, in a concentration–response curve analysis, we found that pelargonidin transactivated AhR with a relative potency, expressed as EC50, of 12 μM (Figure 1B).

**Figure 1.**
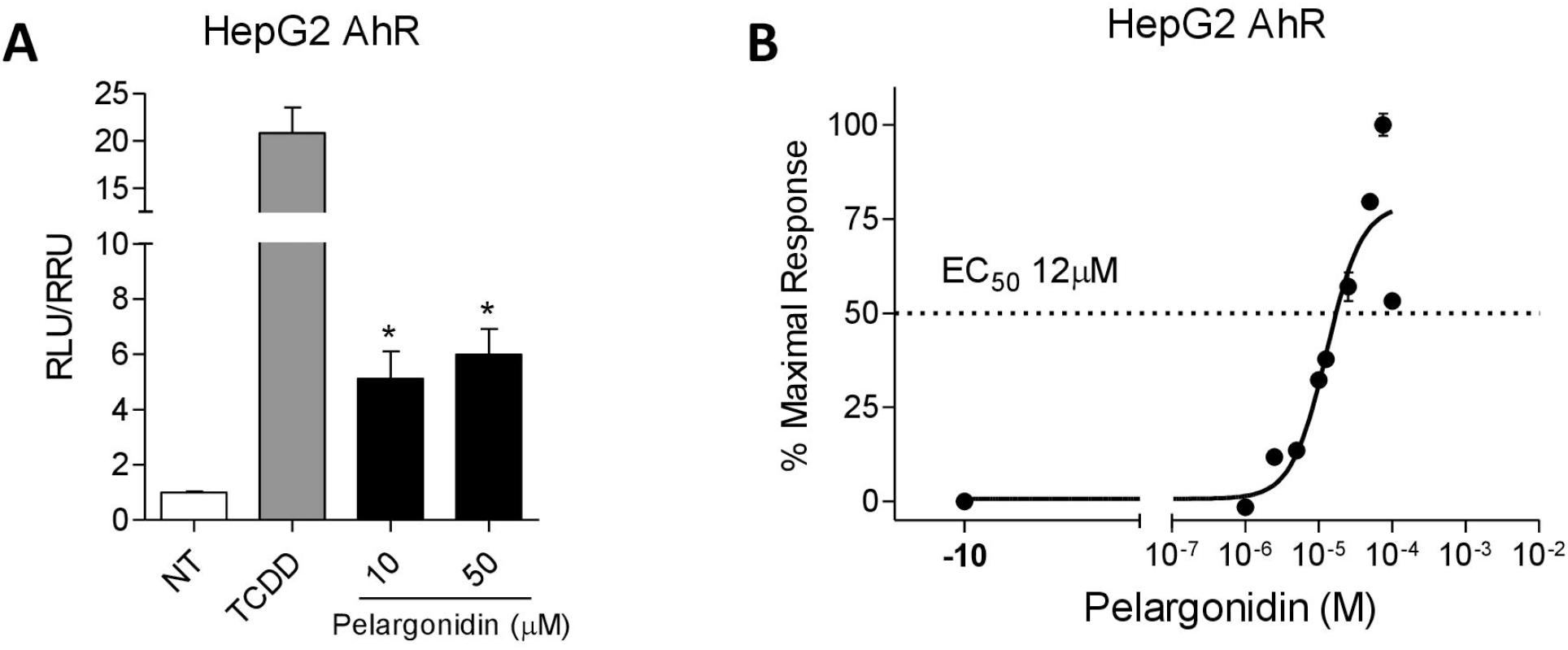
Pelargonidin exerts its effects through the activation of the aryl hydrocarbon receptor (Ahr). (**A**) Fold of induction of luciferase activity in cells transfected with AhR reporter gene and incubated with TCDD (5 nM) or pelargonidin (10-50 μM). (**B**) Dose-response curve of pelargonidin to evaluate AhR activation; cells were stimulated with increasing concentrations of pelargonidin 1 μM to 100 μM. Results are expressed as mean ± standard error. ∗ p < 0.05 versus not treated cells (NT).

### Pelargonidin rescues from intestinal inflammation in TNBS-induced colitis in mice and restore Ace2 expression

We have investigated whether pelargonidin administration attenuates intestinal inflammation in a mouse model of colitis. We first performed a dose finding experiment by administering pelargonidin at a dose of 1, 5 or 10 mg/kg in a mouse model of TNBS-induced colitis (Figure 2). Clinical data and analysis of the macroscopic and microscopic features of the colon showed for doses of 1 and 5 mg/kg of pelargonidin a dose-dependent effect with the lowest dose exerting only mild beneficial effects. However, the 10 mg/kg dose, contrary to expectations, showed intermediate effects between the 1 and 5 mg/kg dose. Because these data suggest that 5 mg/kg of pelargonidin was fully effective in reversing signs of intestinal inflammation we used this dose to further investigate the mechanism of action of pelargonidin on intestinal immunity. For this purpose, we induced a colon inflammation by administering TNBS to Ahr^+/+^ and Ahr^−/−^ mice. As shown previously by us and others, Ahr^−/−^ mice are very susceptible to the TNBS colitis with high mortality rate [26]. In our experimental set we found a mortality rate of 80% in the Ahr knock-out group and therefore it was not possible to further investigate in this strain (data not shown). In wild-type mice, however, the severity of wasting disease and intestinal inflammation induced by TNBS was reversed by treatment with pelargonidin as measured by lower body weight lost, lower CDAI and by assessing the macroscopic and microscopic feature of the colon (Figure 3A-C). In this mouse model of colitis, we found a much greater upregulation of pro-inflammatory cytokine expression than that found in the mouse model of HFD-F (Figure 3E-I); *Il-1β* was up-regulated ≈20 times in the colon of TNBS-treated mice in comparison to ≈2 folds in mice subjected to HFD-F compared to naïve mice (Figure 3G). Furthermore, a five fold increase in *Tnf-α* mRNA was detected in the TNBS model in comparison to mice fed a HFD-F (Figure 3H). These changes were consistent with a very robust increase with in *Ace2 gene* expression (7-fold in comparison to wild type mice). Changes in Ace2 expression associated with robust decrease in *Mas* expression (Figure 3E, F). Administering TNBS mice with pelargonidin reversed this patter by up-regulating the expression of *Tgf-β* and simultaneously decreasing the expression of both *Il-1β* and *Tnf-α* (Figure 3G-I). The anti-inflammatory effect exerted by pelargonidin also resulted in a strong reduction of *Ace2 mRNA* expression. The correlation analysis between *Il-1β*/*Ace2* and *Tnf-α*/*Ace2* confirmed that there was a statistical correlation between the expression of the two pro-inflammatory cytokines and *Ace2* (*Il-1β*/*Ace2 P value = 0.0244; Tnf-α*/*Ace2 P value < 0.0001)*. Again, there was a closer correlation between the expression of *Ace2* and that of *Tnf-α* (*Il-1β*/*Ace2 R squared = 0.1867; Tnf-α*/*Ace2 R squared = 0.7695)*.

**Figure 2.**
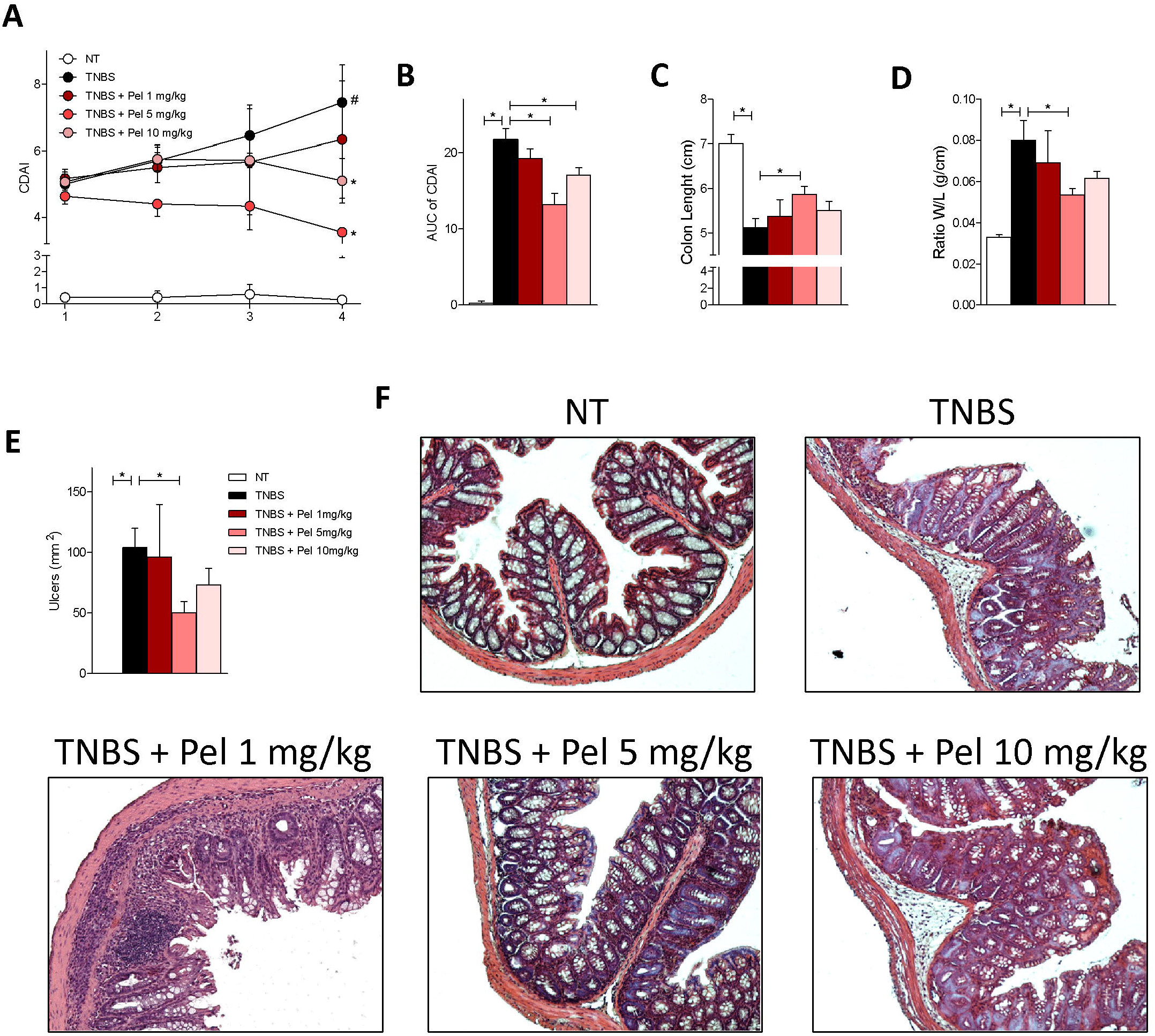
Pelargonidin reduces the severity of TNBS colitis in a dose-dependent manner. Colitis was induced by TNBS. After induction of colitis the mice were treated daily with Pelargonidin (1, 5 or 10 mg/Kg) or vehicle. Disease was monitored by daily evaluation of (**A**) changes in colitis disease activity index (CDAI) and by evaluation of the (**B**) Area Under the Curve (AUC). At the end of the experiment, we evaluated (**C**) colon length (cm) and (**D**) ratio of colon weight/colon length (g/cm). (**E**) Area of ulcers and (**F**) H&E staining of colon sections and Histological Score from each experimental group. Results are expressed as mean ± SEM (n = 5-7); * p <0.05.

**Figure 3.**
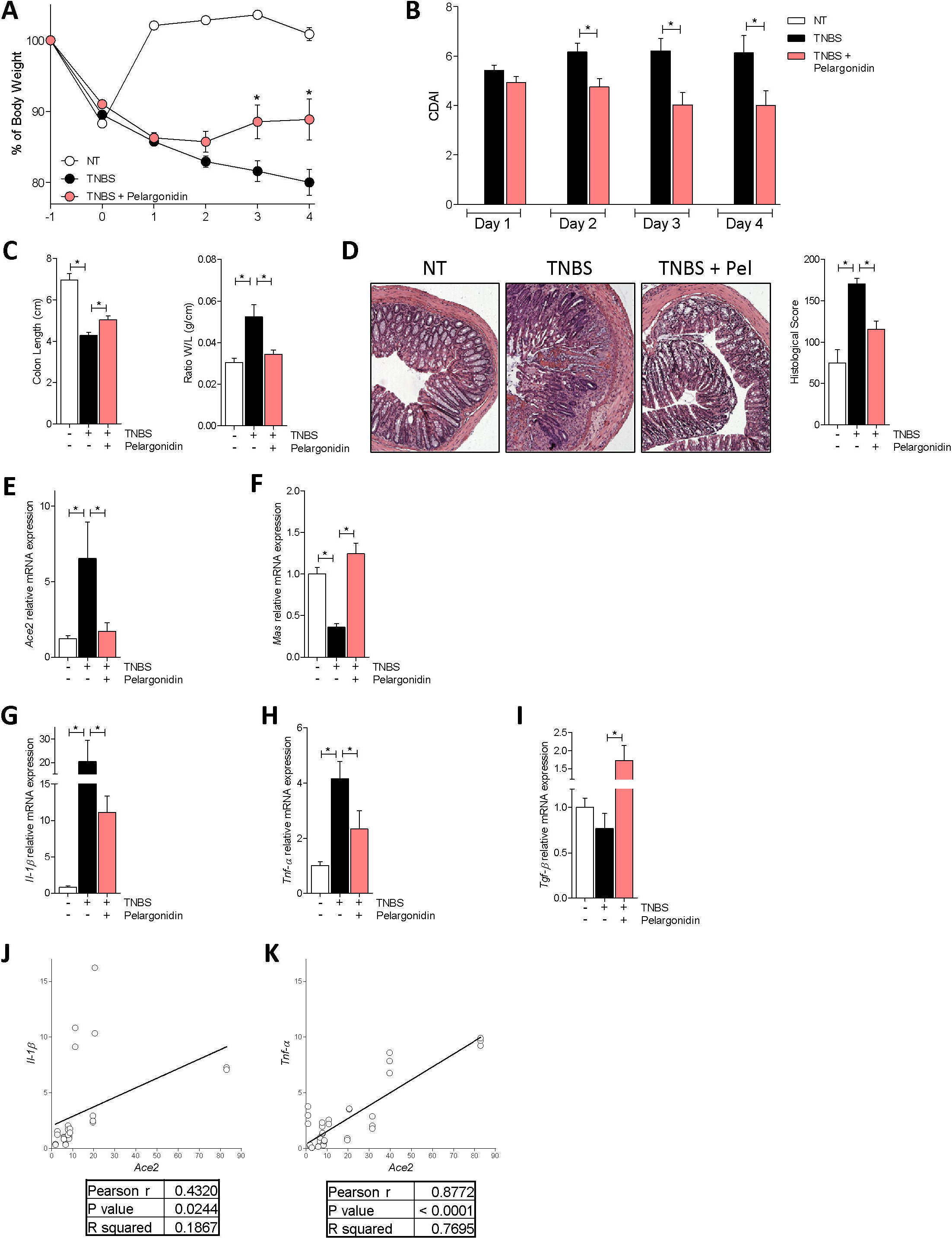
Pelargonidin effect on acute colitis. Colitis was induced by TNBS. After induction of colitis the mice were treated daily with Pelargonidin (5mg/Kg) or vehicle. Disease was monitored by daily evaluation of (**A**) changes in body weight (%), (**B**) colitis disease activity index (CDAI), (**C**) colon length (cm) and ratio of colon weight/colon length (g/cm). (**D**) H&E staining of colon sections and Histological Score from each experimental group. RNA extracted from colon was used to evaluate, by quantitative real-time PCR, the relative mRNA expression of (**E**) *Ace2*, (**F**) *Mas*, (**G**) *Il-1β*, (**H**) *Tnf-α* and (**I**) *Tgf-β*. Values are normalized relative to *Gapdh* mRNA. Correlation graph of *Ace2* mRNA expression and (**J**) *Il-1β* (**K**) *Tnf-α.* Results are expressed as mean ± SEM (n = 7-12); * p <0.05.

Due to the inability to analyze the effects of pelargonidin treatment in the colitis model in Ahr^−/−^ mice for the high mortality, to investigate whether the immunomodulatory effect exerted by pelargonidin was mediated by binding on AHR, we tested the compound on murine macrophages purified from the spleen of Ahr^+/+^ and Ahr^−/−^ mice. Macrophages were subjected to pro-inflammatory stimuli (LPS + IFN-γ) and treated with pelargonidin for 16 h. Stimulation with LPS + IFN-γ induced an increase in the production of pro-inflammatory cytokines (Figure 4A, B) and a reduction in the production of *Tgf-β* (Figure 4C) in macrophages deriving from both genotypes. *Il-6* up-regulation was much higher in Ahr^−/−^ macrophages compared to wild-type (Figure 4E). Treatment with pelargonidin reverted the pro-inflammatory polarization but only in wild-type macrophages, while it had no effect on Ahr^−/−^ macrophages, underlining the role of the AHR receptor in the mechanism of action of pelargonidin.

**Figure 4.**
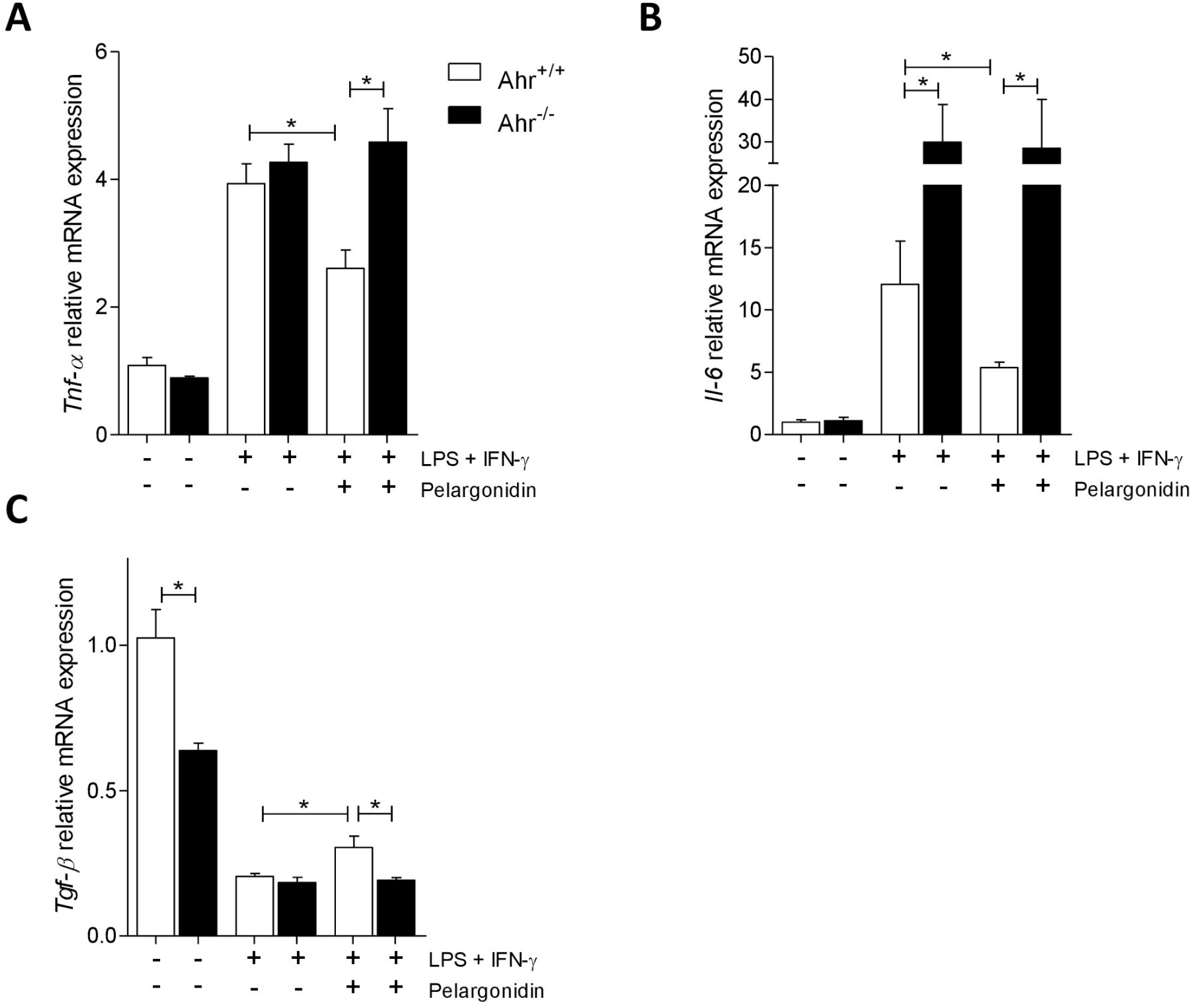
Pelargonidin exert immunomodulatory effects through Ahr. Spleen macrophages purified from AhR^+/+^ and AhR^−/−^ mice were activated in vitro with LPS (5 ng/mL) in combination with IFN-γ (20 ng/mL) alone or plus Pelergonidin (20 μM) for 16h. At the end of stimulation, the relative mRNA expression of pro-inflammatory cytokines (**C**) *Tnf-α* and (**D**) *Il-6*, and anti-inflammatory cytokines (**E**) *Tgf-β*, was evaluated by Real-Time PCR. Values are normalized relative to Gapdh mRNA and are expressed as mean ± SEM (n=6); * p <0.05.

### Pelargonidin protects against colon inflammation induced by HFD-F and down-regulates Ace2 expression

Taking into consideration obesity pandemia and that exposure to high caloric intake promotes a state of intestinal sub-inflammation, and that obesity is a well-defined risk factor for development of severe COVID19, we have first tested the effects of pelargonidin in a mouse model of a high-fat/high-sugar diet [39–41].

In this model, Ahr^+/+^ and Ahr^−/−^ mice were treated with exposed a chronic high caloric [26] intake by feeding them a diet enriched cholesterol and fructose (HFD-F) for 8 weeks. Starting from the second week (day 8) an experimental group of mice for each genotype was treated with pelargonidin daily. As shown in Figure 5, mice exposed to chronic high caloric intake gained significantly more body weight than mice feed a normal chow diet, and they had significantly higher BMI at the end of the study without significant differences between the two genotypes (Figure 5A-C). Treating mice with 5 mg/kg/day pelargonidin for 7 weeks protected against body weight gain and resulted in a lower BMI in comparison to mice feed an HFD-F alone (Figure 5A-C). Instead, the beneficial effects exerted by pelargonidin were completely abrogated in Ahr^−/−^ mice (Figure 5A-C). We have therefore focused our attention on intestinal inflammation. The histological analysis of the colon showed no major morphological abnormalities, although further functional assessment revealed a robust increase of intestinal permeability in the mice exposed to HFD-F (Figure 5D, E). Treating mice with pelargonidin restored the normal colon permeability in wild-type mice, but had no beneficial effects in Ahr−/− mice which exhibited much more severe changes in permeability than in Ahr^+/+^ mice (Figure 5D, E).

**Figure 5.**
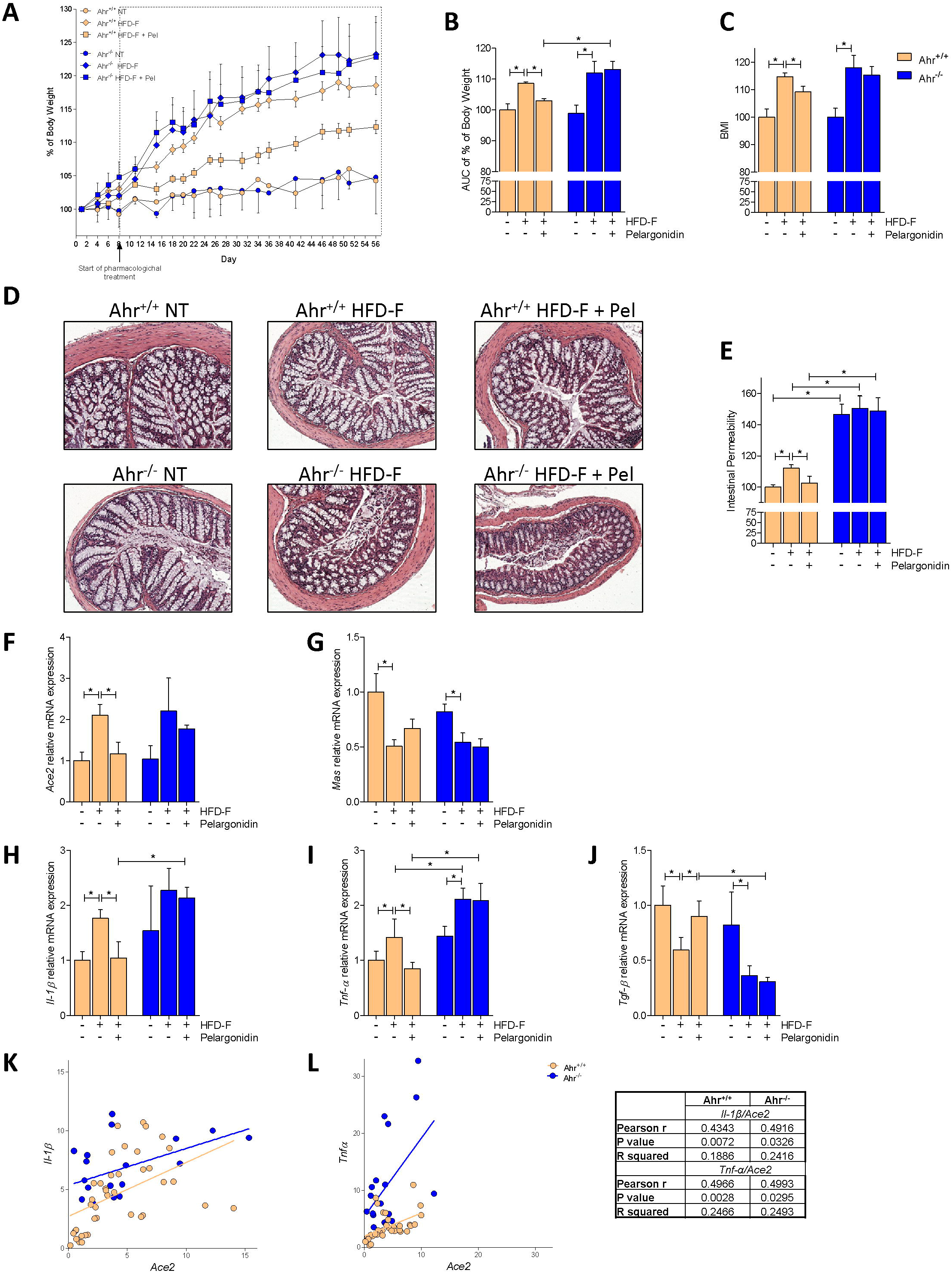
Benefit of Pelargonidin administration in mouse model of NASH is lost in Ahr^−/−^ strains. C57BL/6 male mice, (Ahr^+/+^) and their congenic littermates AhR knock out (Ahr^−/−^) were fed a normal chow diet (NT) or a high fat diet with fructose in water (HFD-F) as described in Material and Methods section. (**A**) Changes in body weight (%) assessed for 56 days. (**B**) Areas under curve (AUC) of body weight expressed in arbitrary units. (**C**) Body Mass Index (BMI) is calculated at the end of the study as the ratio between body weight (g) and body length^2^ (cm^2^). (**D**) Histological sections, performed with H&E staining on colon section of Ahr^+/+^ and Ahr ^−/−^ mice for each experimental group. (**E**) Intestinal permeability was measured after 4 weeks of diet with FITC-dextran administration. At the end of experiment the total RNA extracted from colon was used to evaluate, by quantitative real-time PCR, the relative mRNA expression of (**F**) *Ace2*, (**G**) *Mas*, (**H**) *Il-1β*, (**I**) *Tnf-α*, (**J**) *Tgf-β*. Values are normalized relative to *Gapdh* mRNA. Correlation graph of Ace2 mRNA expression and (**K**) *Il-1β* (**L**) *Tnf-α*. Results are expressed as mean ± SEM (n = 6-10); * p <0.05.

Elevated colonic ACE2 levels have been detected in the colon of IBD patients and are associated with inflammation and severe disease, suggesting a possible correlation between Ace2 expression and colon inflammation in IBD patients [12]. We have therefore investigated the expression of *Ace2* and *Mas* in the colon and compared them with colon profiles of various cytokines (Figure 5F-J). HFD-F increased *Ace2* expression and reduced *Mas* expression in both Ahr^+/+^ and Ahr^−/−^ mice, while increased the colon expression of *Il-1β* and *Tnf-α* and downregulated the *Tgf-β mRNA*. Pelargonidin treatment reversed the inflammatory pattern by down-regulating the expression of both *Il-1β* and *Tnf-α,* but only in Ahr^+/+^ mice. In wild-type mice exposure to the anthocyanin also reduced the expression of Ace2, bringing it back to the value measured in untreated wild-type mice, while exerted only a slight effect on the expression of Mas mRNA (Figure 2F-J). To investigate whether there was also in this mouse models with a mild inflammation in the colon, a direct correlation between the expression of Ace2 and pro-inflammatory cytokines, we performed correlation analyses between *Il-1β* and *Ace2* and between *Tnf-α* and *Ace2.* As shown in Figure 5K and L, the colon levels of the two cytokines were positively correlated with *Ace2* expression in both Ahr^+/+^ and Ahr^−/−^ mice. The fact that the R squared value was greater in the *Tnf-α*/*Ace2* correlation than *Il-1β*/Ace2 confirmed a stronger correlation between the expression of *Ace2* and *Tnf-α*.

### In vitro characterization of pelargonidin effects on intestinal epithelial cells

Because, ACE2 expression in the intestine is highly restricted to epithelial cells, we have then examined whether exposure of Caco-2 cells, a human intestinal epithelial cell line, to TNF-α modulates the expression of ACE2. The results of these experiments demonstrated that exposure to TNF-α promotes the expression of pro-inflammatory cytokines Il-6 and Il-1β by Caco-2 cells (Figure 6B, C) and increased the expression of Ace2 (≈2.5 times) confirming the close correlation between TNF-α and Ace2 (Figure 6A). Administration of pelargonidin reversed this pattern in a concentration-dependent manner (Figure 6A-C).

**Figure 6.**
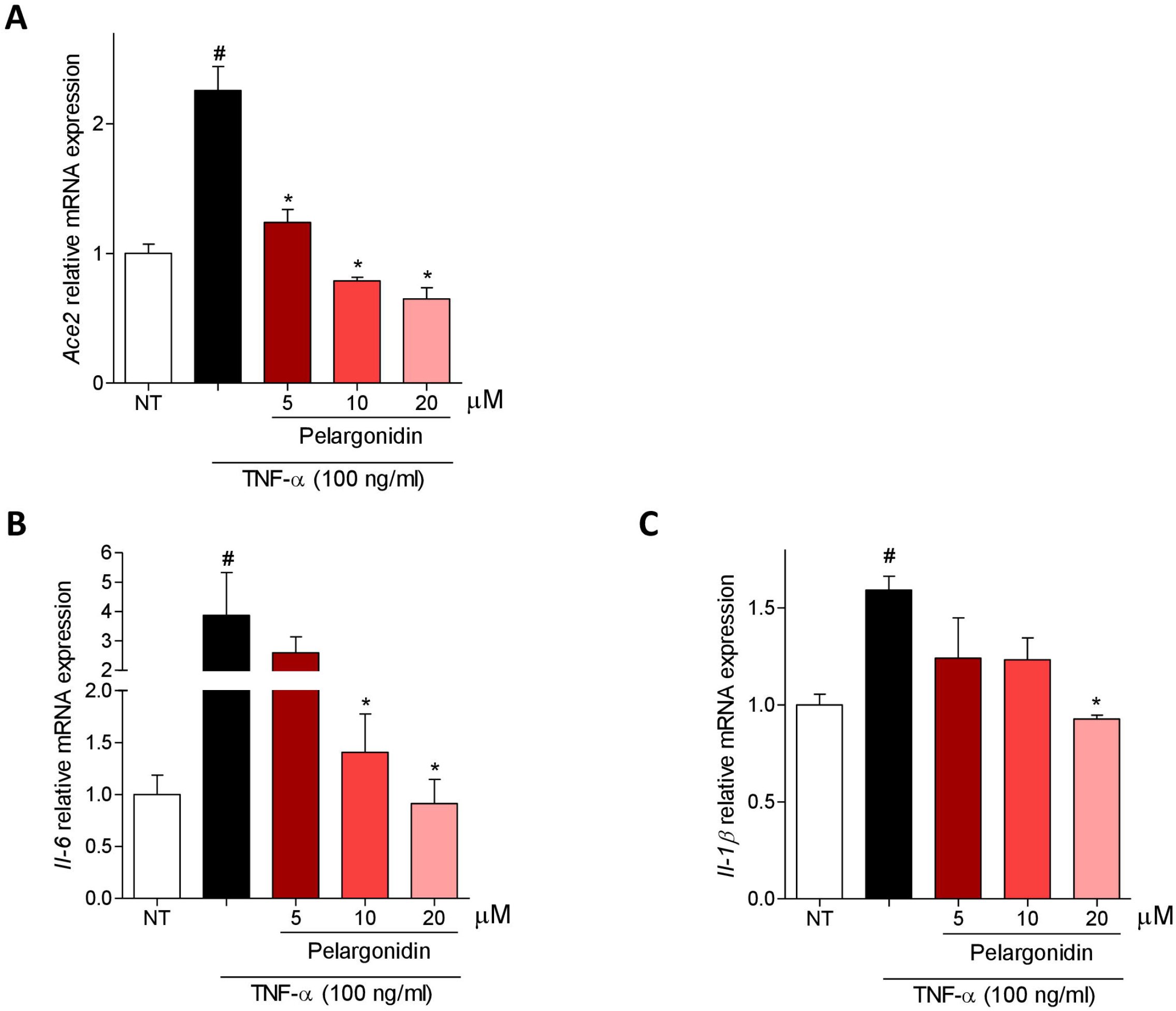
Pelargonidin counteract TNF-α-inflammatory activation on intestinal epithelial cells. Caco-2 cells, a human intestinal epithelial cell line, activated with TNF-α 100 ng/ml for 24 h alone or in combination with pelargonidin (5, 10, and 20 μM). At the end of stimulation, the relative mRNA expression of (A) *Ace2*, (**B**) *Il-6*, and (**C**) *Il-1β*, was evaluated by Real-Time PCR. Values are normalized relative to Gapdh mRNA and are expressed as mean ± SEM (n=5); # NT Vs TNF-α; * TNF-α Vs TNF-α + Pelargonidin; # and * p <0.05.

Since ACE2 represents the host receptor used by the SARS-Cov2 virus to invade cells, and reducing the expression of ACE2 might represent an interesting host defense mechanism against virus invasion. In this context we also investigated whether pelargonidin was able to reduce the binding of the viral Spike protein on ACE2 (Figure 7).

**Figure 7.**
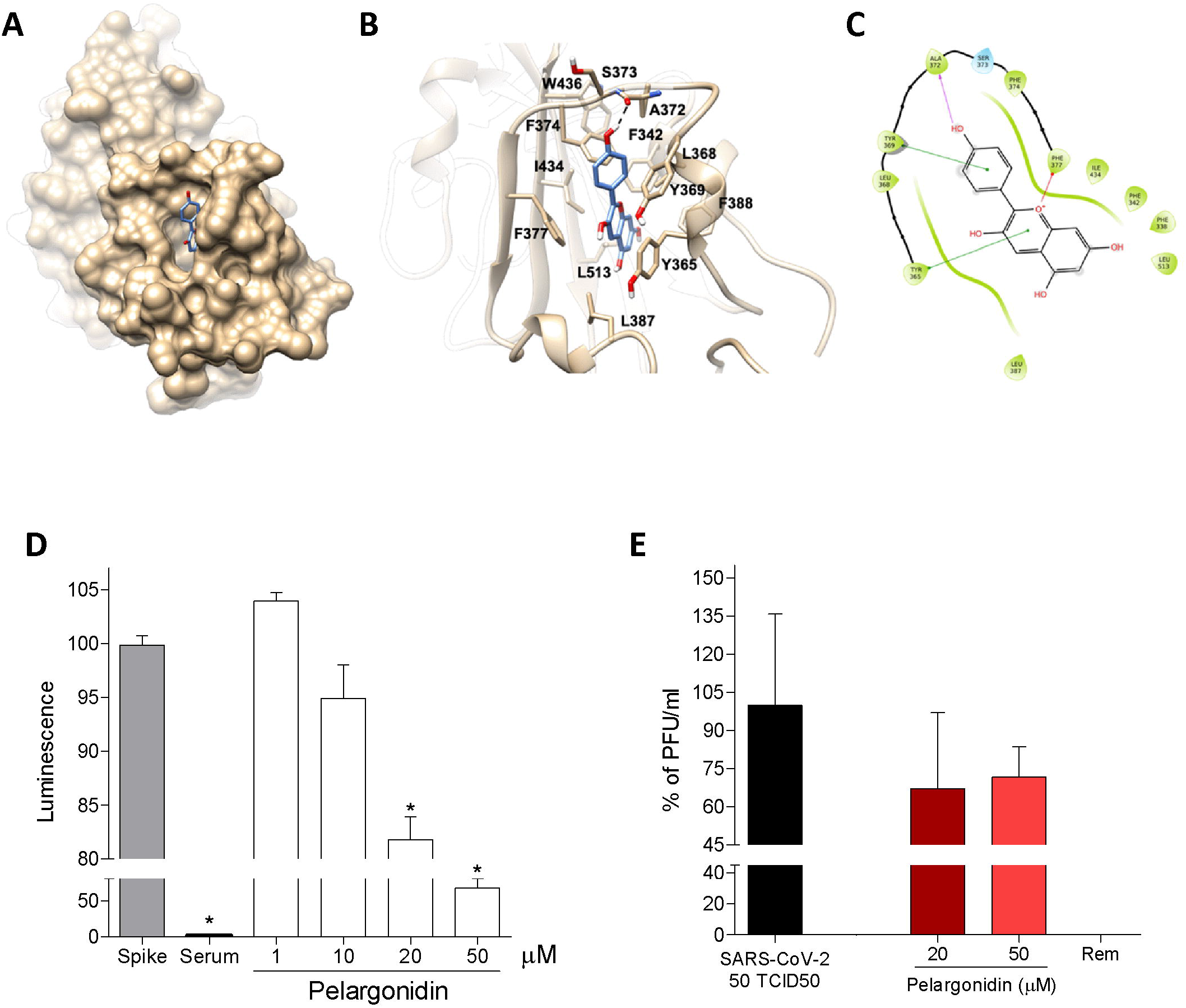
Pelargonidin inhibits binding of the SARS-Cov-2 virus on the host cells. (**A**) Hydrophobic FA binding pocket in a surface representation; (**B**) Cartoon representation of binding mode of pelargonidin to SARS-CoV-2 receptor. The ligand is represented as blue sticks, whereas the interacting residues of the receptor are shown in tan and labelled. Oxygen atoms are depicted in red and nitrogens in blue. The receptors are represented as tan ribbons. Hydrogens are omitted for the sake of clarity; (**C**) Diagram of pelargonidin interaction. (**D**) SARS-CoV-2 Spike binding to immobilized ACE2; Luminescence was measured using a Fluo-Star Omega fluorescent microplate reader. Pelargonidin was tested at different concentration (1, 10, 20 and 50 μM), to evaluate their ability to inhibit the binding of Spike protein (5 nM) to immobilized ACE2, by using the ACE2:SARS-CoV-2 Spike Inhibitor Screening assay Kit. Results are expressed as mean ± SEM (n = 5); * p <0.05. To confirm the validity of the assay used in this study, we tested plasma samples of post COVID-19 patients as a control. (**E**) Virus growth in Vero 6E cells analyzed by plaque assay. Pelargonidin was tested at concentration of 20 and 50 μM. To confirm the validity of the assay used in this study, we tested Remdesivir.

We therefore investigated, through docking calculations, the possibility of pelargonidin to bind several pockets suggested in previous studies: i) the hydrophobic pockets on the β-sheet core of the RBD [42], ii) the fatty acid (FA) pocket identified by Toelzer *et al.* [33, 43]. and finally, iii) the flavonoids binding pocket suggested by Allam *et al*. [44]. All docking calculations were performed using Glide software package. The scores of the best poses highlighted a marked preference of the results obtained in the FA pocket (best docking score −7.7 kcal/mol), with respect to the results obtained in the hydrophobic pockets on the β-sheet core of the RBD (best docking score −5.2 kcal/mol) or in the pocket proposed by Allam (best docking score −5.8 kcal/mol), thus suggesting that pelargonidin binds the FA pocket on the RBD of Spike, as already reported for other highly unsaturated naturally occurring compounds (e.g. Vitamin k, retinoids) [43]. Analysis of the best scored and clustered docking pose in the FA pocket (Figure 7A-C) showed the pelargonidin polyphenolic ring contacting with Leu368, Leu387, Phe388, Phe342 and Ile434. Furthermore, Phe377 makes a π-cation interaction with the oxygen of the ring C, whereas Tyr365 and Tyr369 are engaged in a π-π stacking with the biphenyl ring and with the ring B, respectively. Finally, the binding mode is further stabilized by the H-bond formed between the ring B hydroxyl group and the backbone of Ala372.

Given the results of the docking calculations, we have then investigated whether the pelargonidin impact on the binding of S protein to the ACE2 receptor. For this purpose, a Spike/ACE2 Inhibitor Screening Assay Kit was used.

The results obtained by us showed that pelargonidin was able to reduce binding in a concentration-dependent manner. At the highest tested concentration, equal to 50 μM, we measured a reduction in Spike binding on ACE2 of about 40% (Figure 7D). To confirm this data, we also performed a SARS-CoV-2 virus infection test on Vero E6 cell line, kidney epithelial cells extracted from monkey. In this assay, pelargonidin showed an ability to reduce virus entry, measured by plaque assay, of approximately 30% (Figure 7E).

The data obtained from docking and *in vitro* tests associated with the down-regulation of Ace2 expression exerted by pelargonidin and the anti-inflammatory activity make pelargonidin very interesting in the prevention and treatment of SARS-Cov-2 infection.

## Discussion

Recent data suggest that colon inflammation up-regulates the expression of Ace2. Studies of IBD patients have shown that a direct correlation exists between the severity of colon inflammation and the expression of ACE2 [12, 13, 45].

These data are consistent with previous findings [10, 46] demonstrating that ACE2 gene expression is increased in human bronchial epithelial cells infected by SARS-CoV-2 as a response to inflammatory cytokine stimulation including interferon IFN-γ. In the colon of patients with active IBD several pro-inflammatory cytokines like IFN-γ, TNFα, IL-1, and IL-6 are up-regulated [47]. In line with these data, we wanted to investigate the correlation between pro-inflammatory cytokine expression and Ace2 expression in the colon in a mouse models of IBD and mouse model of high caloric intake. Obesity is a clinical condition with a high prevalence in the most industrialized countries and it is now known that it involves a state of sub-inflammation in the gastrointestinal tract [48, 49] and it is a well-defined risk factor for develop severe COVID19.

ACE2 represents the host receptor used by the SARS-Cov2 virus to invade cells. Two recent studies [50, 51] have shown that the virus might infect the human gut, and therefore the modulation of Ace2 expression in the gastrointestinal tract represents an interesting research area. The data we have obtained in the two mouse models of intestinal inflammation strongly support the notion that upregulation of ACE2 in colon is due to the induction of state of inflammation characterized by strong induction of expression of pro-inflammatory cytokines. Importantly, in contrast to ACE2 the expression of the Ang 1-7 receptor Mas, was markedly downregulated in these models (Figure 3 and 5). In the TNBS model the levels of colon cytokine were increased by 5-20 folds and the expression of ACE2 mRNA was induced by ≈7-fold in comparison to naive mice (Figure 3E). The state of mild inflammation present in the colon of mice fed with HFD-F induces an increase in the expression of Il-1β and Tnf-α by 2 times, while the expression of Tgf-β was markedly reduced (Figure 5H-J). Moreover, in the colon of animals subjected to a high-calorie diet, the expression of ACE2 doubled while that of MAS was halved when compared to that in mice fed a chow-diet (Figure 5F, G). These data establish a direct correlation between the severity of inflammation and ACE2 gene transcription.

In both mouse models we have found that values of R squared indicated a close correlation between the expression of Ace2 and the pro-inflammatory cytokine Tnf-α. These data are in agreement with the reducing effect on ACE2 expression in the colon of IBD patients exerted by therapy with anti-TNF-α [12, 52]. Confirmation of the direct regulation of ACE2 expression exerted by TNF-α was given by the in vitro experiment on human intestinal epithelial cell line (Caco-2). Indeed, the data obtained by us showed that the exposure of intestinal epithelial cells to TNF-α is able to up-regulate the expression of ACE2 (Figure 6A).

Similarly, to other flavonoids, pelargonidin, a common constituent of many red plant foods, including berries, grapes or cabbage, have been used in the traditional medicine in the treatment of a number of human alignments [53, 54]. Pelargonidin display a several biological activities, the most prominent being those related to their anti-oxidant effects. We have confirmed in this study that pelargonidin exerts anti-inflammatory effects in intestines in mouse models of IBD and obesity through the interaction with the AhR. Treatment with pelargonidin reduced the expression of pro-inflammatory cytokines in the colon in both models of intestinal inflammation, alleviating the disease (Figure 2, 3 and 5). Furthermore, treatment with pelargonidin restored ACE2 expression to the level measured in the colon of naive mice (Figure 3E and Figure 5F). The beneficial effects exerted by the compound were abrogated in the absence of AhR confirming the functional relevance of the receptor in mediating the anti-inflammatory activities of this flavonoid (Figure 4, 5). In vitro studies also confirmed that pelargonidin might directly inhibit ACE2 induction caused by exposure of human intestinal epithelial cell line to the TNF- α (Figure 6).

The regulatory effects of pelargonidin on the intestinal ACE2 might be of relevance in the context of SARS-CoV2 infection, since high levels of ACE2 are compatible with an increased risk of SARS-CoV2 uptake by the inflamed colon [12]. Importantly, in addition to ACE2 regulation, we have shown that beneficial effects of pelargonidin might extend to a direct regulation of SARS-CoV2/ACE2 interaction. Indeed, we found that pelargonidin not only reduces the interaction between ACE2 and the Spike protein of the SARS-Cov-2 but also reduces the ability of the SARS-CoV-2 to infect Vero E6 cell line, a monkey kidney epithelial cell line used to investigate the SARs-CoV2 replication (Figure 7). We have investigated the molecular mechanisms involved in preventing the binding of SARS-CoV2 virus to its receptor identifying the fatty acid binding pocket in the RBD as the putative binding site of pelargonidin.

In summary (Figure 8) we shown that the natural flavonoid, pelargonidin, reverses ACE2 induction in rodent models of inflammation and this effect is mediated by AhR. In addition, the pelargonidins might reduce SARS-CoV2/ACE2 interaction affecting the virus uptake and replication. These data highlight the potential of natural agents in reducing inflammation.

**Figure 8.**
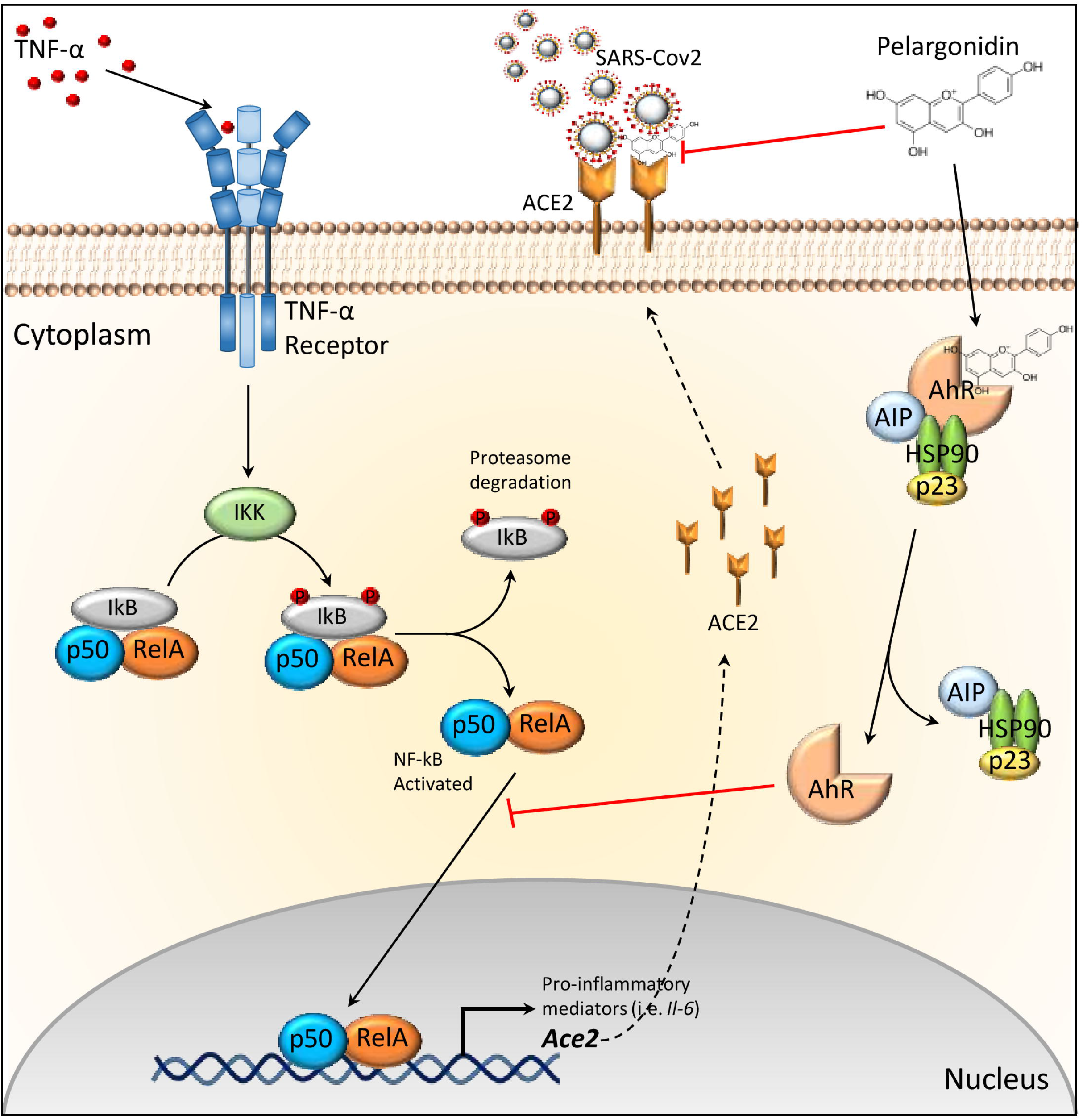
Pelargonidin down-regulates Ace2 expression and inhibits binding of the SARS-Cov-2 virus on the host cell ACE2 receptor. Inflammatory stimuli lead to the release of TNF-α which binds to its receptor present on epithelial cells. The activation of TNF-α receptor induces an increase in IkB phosphorylation by IKK, which is then degraded releasing the p50/RelA dimer that migrates to the nucleus where activates the transcription of several genes including *Il-6* and *Ace2*. Pelargonidin exerts a protective effect with an Ahr-dependent and an Ahr-independent mechanism. The compound activates the Ahr receptor which blocks NF-kB translocation into the nucleus by sequestering the RelA subunit and thereby inhibiting the expression of *Ace2*. Pelargonidin also directly inhibits the binding of the SARS-Cov-2 virus to the ACE2 receptor. This double action, together with the anti-inflammatory activity, inhibits the entry of the virus into cells.

